# LiaS-dependent activation of the MadR regulon enables cross-talk between *Enterococcus faecalis* cell envelope defense systems

**DOI:** 10.1101/2025.06.19.660511

**Authors:** Kara S. Hood, Samie A. Rizvi, Guillermo A. Hoppe-Elsholz, Ana P. Streling, Diana Panesso, Shivendra Pratap, Yousif Shamoo, Cesar A. Arias, William R. Miller

## Abstract

Enterococci are gastrointestinal commensals that must defend their cell envelope against antimicrobial peptides derived from the host and other members of the microbiota. The signaling systems LiaFSR and MadRS are pivotal for survival in the presence of antimicrobial peptides and antimicrobial peptide-like antibiotics such as daptomycin. Both systems possess a signaling histidine kinase (LiaS, MadS) and cognate response regulator (LiaR, MadR) that activate transcription of distinct sets of effector genes. Using isogenic deletion strains, we noted differences in daptomycin minimum inhibitory concentration (MIC) between the laboratory strain *E. faecalis* OG1RF (1.5 µg/mL), OG1RFΔ*madR* (0.38 µg/mL), and OG1RFΔ*madS* (4 µg/mL). Transcriptional analysis of the MadR regulon showed a daptomycin-dependent increase in *madG, madL*, and *dltA* gene expression in the OG1RFΔ*madS* background, suggesting activation of the LiaFSR system may provide a cross-regulatory role. Deletion of the *liaS* gene in OG117Δ*madS* was associated with a significant decrease in daptomycin MIC and loss of *madG* expression, one of the most differentially expressed genes on activation of the MadR regulon. Using microscale thermophoresis, LiaS showed a similar binding affinity to both LiaR (K_d_ 2.42 µM) and MadR (K_d_ 5.02 µM), while MadS showed higher affinity for its cognate regulator MadR (K_d_ 8.08 µM) than for LiaR (K_d_ 25.7 µM). Taken together, our findings indicate that the MadR regulon can be expressed independent of MadS-induced signaling, likely through cross-talk between LiaS and MadR. Thus, enterococci have evolved an interconnected network of cell envelope signaling that permits bacterial survival in the presence of antibiotics and antimicrobial peptides.

## INTRODUCTION

Enterococci are leading causes of healthcare associated infections and display intrinsic resistance to a variety of frequently used antibiotics such as cephalosporins [1]. They are commensals of the gastrointestinal tract microbiota and must defend the cell envelope from lantibiotics, bacteriocins, and host derived peptides secreted into the environment [2]. The enterococcal cell envelope stress response has evolved to provide an overlapping network of signaling systems and effector proteins to meet this threat [3-6]. Specialization in these complementary pathways allows targeted protection of the synthesis and maintenance of cell wall peptidoglycan as well as the phospholipid membrane.

The response to antimicrobial peptides in *Enterococcus faecalis* is primarily mediated by two signaling pathways. The first is LiaFSR (for lipid-II interfering antibiotic response), a three-component system composed of a histidine kinase (LiaS), a response regulator (LiaR), and a putative activator (LiaF) [3]. LiaR-mediated signaling is essential for resistance to daptomycin and the host-derived peptide LL-37, as *liaR* knockout strains exhibit a hypersusceptible phenotype [7]. LiaR regulates a three gene operon, *liaXYZ*, that orchestrates resistance via membrane lipid remodeling through a secreted sensor LiaX and two transmembrane proteins, LiaY and LiaZ [8, 9]. The second pathway is the MadRS (for membrane antimicrobial peptide defense) system, which was originally characterized in the context of bacitracin resistance [10]. In this two-component system, phosphorylation of the MadR response regulator by the MadS histidine kinase depends on the activity of a Bce-like integral membrane transporter (MadAB) which removes bacitracin from its undecaprenyl pyrophosphate target [11, 12]. Thus, activation of the system relies on the “flux” of antibiotic through MadAB [13]. Downstream, MadR controls the expression of several effector systems, including MadLM (responsible for bacitracin resistance), MadEFG (responsible for resistance to LL-37 and human β-defensin 3), MprF2 (which synthesizes lysyl-phosphatidylglycerol), and the *dlt* operon (which mediates D-alanylation of teichoic acids) [14]. These downstream systems either directly remove antibiotics or antimicrobial peptides from their site of action (MadLM and MadEFG), or alter cell surface charge (MprF, Dlt) to provide electrostatic repulsion of cationic peptides.

Prior studies identified a LiaR binding site upstream of *madRS* and demonstrated that transcription of the *madRS* genes is upregulated in a LiaR-dependent manner [15]. This upregulation is thought to prime the cell for defense against membrane-active peptides. Our previous work further showed that mutations which activate MadS can confer daptomycin resistance via expression of the *dlt* operon [14]. However, several important questions remain. Since antibiotic flux through MadAB is required for MadS signaling, it is unclear how the MadR regulon plays a role in resistance to antibiotics not directly recognized by MadAB. In the wild-type system, increased production of MadRS under LiaR induction upon exposure to daptomycin would not be expected to result in a transcriptional response from the genes of the MadR regulon, as daptomycin is not a substrate for the MadAB flux sensor. The aim of this study was to investigate the link between the LiaFSR and MadRS defense systems and dissect the mechanism by which daptomycin activates the MadR regulon in the absence of its cognate signaling members (MadAB-MadS).

## RESULTS AND DISCUSSION

### The *madS* gene is not required for the *E. faecalis* OG1RF response to daptomycin and bacitracin

To characterize the contributions of the histidine kinase MadS and its cognate response regulator MadR to the daptomycin response, we utilized single gene isogenic deletion strains in the laboratory *E. faecalis* OG1RF background. Consistent with previous results, deletion of *liaR* and *madR* resulted in a decrease in daptomycin MIC, from 1.5 µg/mL in the wild-type strain to 0.047 µg/mL and 0.38 µg/mL, respectively (**Table 1**). Similar reductions were seen in bacitracin MIC, from 32 µg/mL in wild-type OG1RF, to 4 µg/mL in OG1RFΔ*liaR* and 12 µg/mL in OG1RFΔ*madR*. Interestingly, deletion of *madS* from OG1RF resulted in an increase in daptomycin MIC from 1.5 to 4 µg/mL and no change in the bacitracin MIC.

**Table 1.** Strains and antimicrobial minimum inhibitory concentrations (MIC, µg/mL).

| Strain | DAP | BAC |
| --- | --- | --- |
| OG1RF | 1.5 | 32 |
| OG1RFΔ*liaR* | 0.047 | 4 |
| OG1RFΔ*madR* | 0.38 | 12 |
| OG1RFΔ*madS* | 4 | 32 |
| OG117 | 1.5 | 32 |
| OG117Δ*liaS* | 0.125 | 8 |
| OG117Δ*madS* | 4 | 32 |
| OG117Δ*madS*Δ*liaS* | 0.094 | 2 |

Given the surprising increase in daptomycin MIC and stable bacitracin phenotype with the loss of signaling through MadS, we performed qRT-PCR to evaluate for transcriptional changes in genes belonging to the MadR regulon (**Figure 1**). In the strains grown in the absence of antibiotics, we saw a significant decrease in expression of *madG* and *madL* in both the OG1RFΔ*madR* and OG1RFΔ*madS* strains, as compared to wild-type OG1RF. When exposed to a sub-inhibitory concentration of daptomycin, there was no transcriptional activation of *madG, madL*, or *dltA* in the OG1RFΔ*madR* strain, as expected. However, in OG1RFΔ*madS* expression of all three genes was similar to that of wild-type OG1RF. Thus, on daptomycin exposure *E. faecalis* OG1RF was able to activate the MadR regulon in a MadS-independent manner.

**Figure 1.**
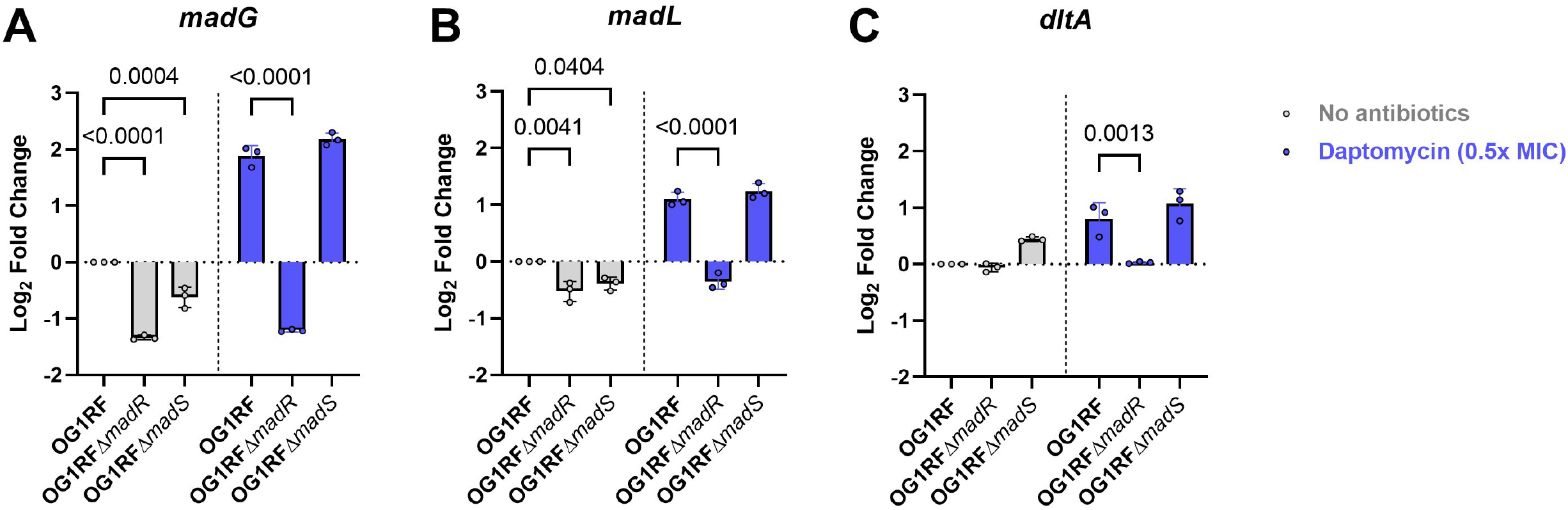
Daptomycin induces activation of the MadR regulon in a MadS independent manner. Fold-change in gene expression for the strains *E. faecalis* OG1RF, OG1RFΔ*madR*, and OG1RFΔ*madS* as measured by qRT-PCR for **A)** *madG*, **B)** *madL*, and **C)** *dltA*, in TSB without antibiotics (grey bars) or TSB plus 50 mg/L calcium chloride and daptomycin at 0.5 x the MIC (blue bars). Individual biological replicates are shown as small circles, error bars represent standard deviation, and significant p-values are indicated above the respective comparisons.

The unexpected results of activation of the MadR regulon in the absence of MadS suggested an alternative mechanism of MadR phosphorylation that bypasses the canonical MadAB-MadS mediated pathway. While it was previously shown that LiaR leads to an increase in expression of MadRS (termed SapRS) [15], it is not known how antibiotics such as daptomycin, which are not substrates for the MadAB flux sensor required for MadS function, would lead to the expression of effector systems important for resistance (such as *mprF* and the *dlt* operon). Since the LiaFSR system induces expression of MadRS and plays a pivotal role in the enterococcal response to daptomycin, we hypothesized that LiaS may play a direct role in the activation of the MadR regulon.

### Deletion of *liaS* abolishes activation of the MadR regulon in the absence of *madS*

To test the above hypothesis, we constructed isogenic deletions of *liaS, madS*, and a *madS/liaS* double deletion using the *E. faecalis* strain OG117. Deletion of *liaS* alone was sufficient to reduce the daptomycin MIC to 0.125 µg/mL and bacitracin MIC to 8 µg/mL, while OG117Δ*madS* displayed a daptomycin MIC of 4 µg/mL and a bacitracin MIC of 32 µg/mL (**Table 1**). Moreover, in the OG117Δ*madS*Δ*liaS* strain, both the daptomycin MIC (0.094 µg/mL) and bacitracin MIC (2 µg/mL) were reduced to a level similar to the *liaR* deletion strain.

We next evaluated the expression of *madR* and *liaX* (both transcriptionally regulated by LiaR but not MadR), and *madG* (transcriptionally regulated by MadR but not LiaR) to better characterize the interplay between LiaFSR and MadRS systems across different genetic backgrounds. In wild-type OG117, both bacitracin and daptomycin lead to small increases in the transcription of *madR*, by 1.1-fold and 1.9-fold respectively (**Figure 2A**). As expected, deletion of *liaS* abolished the antibiotic-dependent activation of genes under the control of LiaR, with *madR* and *liaX* expressed at a level comparable to the basal transcription of uninduced wild-type OG117 (**Figure 2A and 2C**). Consistent with the MIC data, the OG117Δ*madS* strain showed persistent activation of *madG* expression under both bacitracin (2.2-fold) and daptomycin (1.3-fold) exposure, along with preserved expression of *liaX* (**Figure 2B and 2C**). In the OG117Δ*madS*Δ*liaS* strain, however, there was a significant decrease in the transcription of *madG* even under antibiotic induction, by -4.8-fold under bacitracin exposure and -3.3-fold under daptomycin exposure (**Figure 2B**).

**Figure 2.**
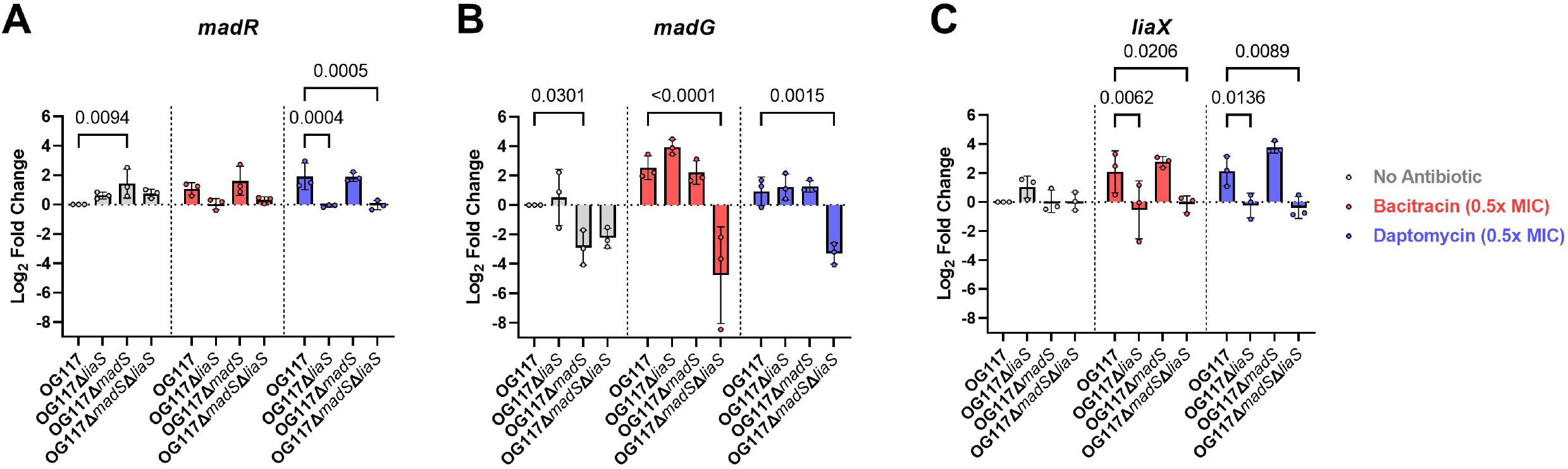
Differential expression of the MadR and LiaR regulons in histidine kinase deletion strains. Fold-change in gene expression for the strains *E. faecalis* OG117, OG117Δ*liaS*, OG117Δ*madS*, and OG117Δ*madS*Δ*liaS*, as measured by qRT-PCR for **A)** *madR* (under control of LiaR), **B)** *madG* (under control of MadR), and **C)** *liaX* (under control of LiaR), in TSB without antibiotics (grey bars), TSB plus bacitracin zinc at 0.5 x the MIC (red bars), or TSB plus 50 mg/L calcium chloride and daptomycin at 0.5 x the MIC (blue bars). Individual biological replicates are shown as small circles, error bars represent standard deviation, and significant p-values are indicated above the respective comparisons.

In the absence of antibiotic stress, loss of the MadS histidine kinase results in decreased expression of MadR-regulated genes. However, under antimicrobial stress induced by daptomycin and bacitracin both of which activate the LiaFSR response, we found increases in transcription of downstream genes in the MadR regulon in the absence of MadS. This activation required the presence of LiaS, as the double knockout strain lacking both *madS* and *liaS* showed no activation of either system.

### Characterization of LiaS and MadR interaction using microscale thermophoresis

The results of the antimicrobial susceptibility tests and transcriptional analyses suggest that LiaS can substitute for MadS in activating the MadR regulon. Therefore, we investigated the binding kinetics of each histidine kinase with both response regulators using microscale thermophoresis (**Figure 3**). LiaS was found to have a similar binding affinity for both its cognate response regulator LiaR (K_d_ 2.42 µM) and the non-cognate regulator MadR (K_d_ 5.02 µM). MadS, however, displayed a higher affinity for the cognate regulator MadR (K_d_ 8.08 µM) than the LiaR response regulator (K_d_ 25.7 µM). Neither kinase showed any binding with lysozyme, which was used as a negative control. Thus, LiaS associates *in vitro* with both LiaR and MadR, and this binding affinity mirrors the activation of both regulons observed in the transcriptional data.

**Figure 3.**
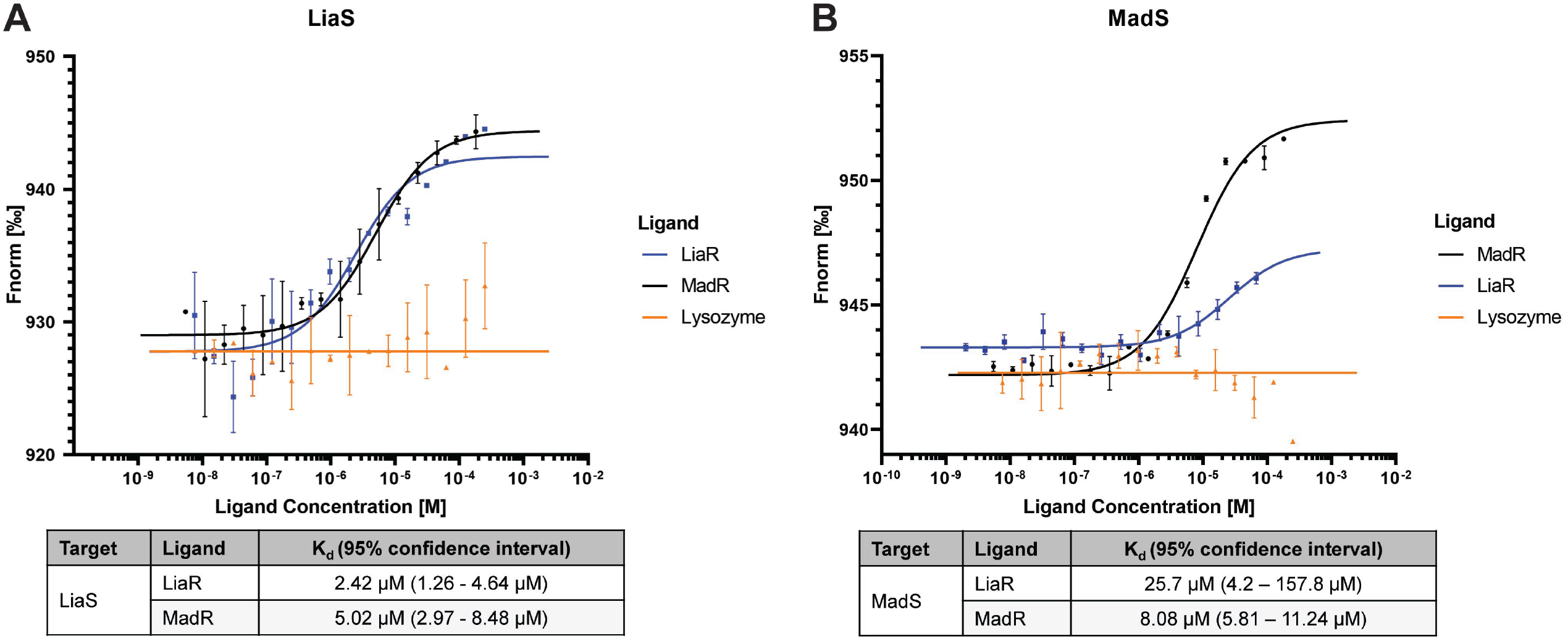
Binding affinity of histidine kinase/response regulator pairs using microscale thermophoresis (MST). Binding curves showing ligand concentration on the X-axis and normalized fluorescence on the Y-axis. Labeled targets were **A)** LiaS and **B)** MadS at fixed 20 nM concentration. Dissociation constants (K_d_) for histidine kinase-response regulator pairs are given below the corresponding graph. Lysozyme was used as a negative control.

An interesting observation is the directionality of cross-regulation, as we did not observe any activation of the LiaR regulon in strains lacking LiaS. First, since expression of the *madRS* genes are under LiaFSR control, basal levels of MadS and MadR may not be sufficient to generate a transcriptional response. Second, our microscale thermophoresis data suggest that the affinity of MadS is greater for the cognate regulator MadR, with a lower affinity for LiaR. This contrasts with LiaS, which displayed roughly equivalent K_d_ for each response regulator. Thus, there appears to be a layered response, with the more general LiaFSR system leading to cross-activation of the MadRS system, but not the reverse (direct activation of LiaFSR by MadS).

There are several potential benefits to the organization of dual activation of signal transduction by a non-cognate histidine kinase (**Figure 4**). LiaFSR is a global sensor of cell envelope stress and membrane perturbations. Thus, rapid activation of a general membrane defense including remodeling of lipid species via LiaY-Cls and alteration of membrane charge would be beneficial for cell survival [8, 9]. While this could be accomplished under a single LiaR regulon, the presence of two separate response regulators would allow a more nuanced response to peptides and lantibiotics with multiple mechanisms of action. For molecules acting against undecaprenyl pyrophosphate, the canonical substrates of Bce-like transporters, the increase in flux via MadAB would lead to a specific upregulation of targeted effectors of resistance, restoring membrane homeostasis and more quickly quenching the stress response [16]. In the presence of molecules which are not MadAB substrates, such as daptomycin, cross signaling via LiaS would ensure continued expression of genes important for modulating surface charge, while LiaY mediated diversion of anionic phospholipids directs the primary mechanism of daptomycin defense in *E. faecalis* [9, 17]. This split signaling could help mitigate the fitness cost of such a broad response as compared to a single activator.

**Figure 4.**
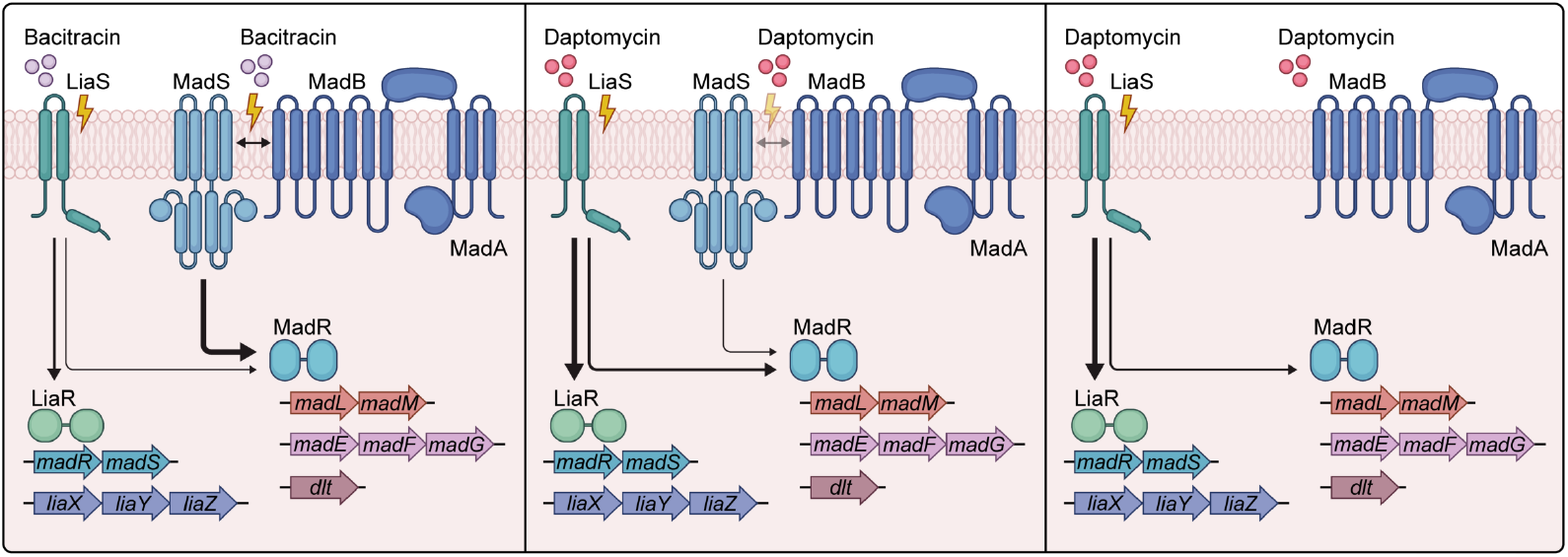
Cross-regulation of the MadRS system by LiaS. Left panel) Bacitracin activates both the general membrane stress response (via LiaFSR, green) and the MadR regulon (via MadAB-MadS, blue) to express downstream effector genes. **Middle panel)** Daptomycin activates the LiaFSR response but is not an efficient substrate for the MadAB-MadS flux sensor. However, LiaS dependent activation of MadR ensures transcription of *dlt* and *mprF* (not pictured). **Right panel)** In the absence of MadS, activation of the LiaFSR stress response directly acts on MadR, bypassing the need for MadS signaling.

In summary we show that the MadR regulon can be activated in a LiaS-dependent manner in *E. faecalis* OG1RF. This cross-activation reveals a tiered response to membrane active antibiotics such as daptomycin and bacitracin, suggesting that the inactivation of the LiaFSR system may disarm two critical cell envelope stress response signaling networks at once. Further studies are warranted to explore this strategy in clinical isolates of *E. faecalis*, and to understand if cross regulation of these systems plays a role in the membrane defense of *E. faecium*.

## MATERIALS AND METHODS

### Bacterial strains and susceptibility testing

Enterococci were cultured on brain heart infusion (BHI) agar or in BHI broth (Oxoid, ThermoFisher) with rolling agitation at 37° C unless otherwise specified. The strains *E. faecalis* OG1RFΔ*liaR* and OG1RFΔ*madR* were previously published [4, 7]. OG1RFΔ*madS*, OG117Δ*madS*, OG117Δ*liaS*, and OG117Δ*madS*Δ*liaS* were generated for this work using the pHOU1 plasmid (for the *madS* deletion) or the pCE plasmid and CRISPR-*cas9* system (for the *liaS* deletion) as previously described [4, 14, 18]. Primers are provided in **Supplemental Table 1**. Minimum inhibitory concentrations (MICs) were performed using gradient diffusion strips of daptomycin (Etest, BioMérieux) and bacitracin (MTS, Liofilchem) on Mueller Hinton agar (Becton Dickinson, ThermoScientific).

### Quantitative reverse-transcription PCR (qRT-PCR)

The qRT-PCR assays were performed as described previously [14]. Briefly, strains were grown overnight in tryptic soy broth (TSB, Becton Dickinson, ThermoScientific), which was supplemented with 50 mg/L of calcium chloride for strains exposed to daptomycin or their controls. All strains were then diluted in fresh TSB or TSB plus calcium chloride as appropriate, and daptomycin or bacitracin-zinc were added at one-half the MIC for the respective strain. Isolates were grown until mid-logarithmic phase (OD_600_ of 0.5-0.7), pelleted, then flash frozen until RNA isolation. Total RNA was isolated with the PureLink RNA kit (Invitrogen), removal of genomic DNA was performed using TurboDNase (Ambion), and synthesis of cDNA was carried out with SuperScript II reverse transcriptase (Invitrogen). Expression of target genes was measured with SYBR green on a CFX96 Touch real-time PCR detection system (Bio-Rad). Primer sequences used for detection are provided in **Supplemental Table 1**. Gene expression was normalized to *gyrB*, and fold-change was calculated using the Pfaffl method. Differences in expression were compared to the wild-type OG1RF or OG117 *E. faecalis*, as appropriate. Statistical significance was defined as a p-value of <0.05 determined using two-way ANOVA with Tukey’s test for multiple comparisons (Prism, GraphPad). Each experiment was performed in three biological replicates, with technical triplicates for each assay.

### Protein expression and purification

Full-length LiaSand, LiaR, with N-terminal His6 tags were expressed from the inducible pET28a vector in *E. coli* Lemo-21 (NEB),while *MadS* and *MadR* were expressed under the same conditions in *E. coli* BL21 (DE3). The cultures were induced at OD_600_ ∼0.6 with 400 µM IPTG at 37° C for 4 hours before harvest. Subsequently, cells were pelleted and washed once before storing at -80° C. To purify LiaS, cell pellets were thawed on ice before resuspending in lysis buffer (100 mM sodium phosphate pH 8.0, 200 mM sodium chloride, 5% glycerol, 1 mM PMSF and EDTA-free protease inhibitor cocktail [Pierce]). Cells were lysed at 1500 psi by French Press before crude lysate separation. Membranes were collected by ultracentrifugation at 150,000x g at 4° C for 1 hour and resuspended in lysis buffer with 1% n-dodecyl-B-D-maltoside (DDM, w/v) detergent. Membranes were solubilized for 1 hour at 4° C before separating the insolubilized fraction by centrifugation at 30,000x g at 4° C for 30 minutes. The soluble fraction was incubated rotating overnight at 4° C with TALON Cobalt resin primed in lysis buffer plus 1 mM imidazole. After loading into a column, the resin was washed 3 times in increasing imidazole concentrations (5 mM, 25 mM, 45 mM) before being eluted with 300 mM imidazole in 1.5 mL fractions. Fractions were subject to SDS-PAGE to identify those containing LiaS, which were then pooled and concentrated to <3 mL with 10K MWCO Amicon ultracentrifugal filters. The concentrated sample was loaded on a BioScale Mini Bio-Gel P6 Desalting column (BioRad) primed with LiaS storage buffer (100 mM sodium phosphate pH 8.0, 100 mM sodium chloride, 10% glycerol, 1 mM PMSF, and 0.01% DDM (w/v)). Elution fractions were further concentrated to <500 µL as described above and injected onto a Superdex 200 Increase (10/300) size exclusion column (Cytiva). LiaS fractions were identified with SDS-PAGE, pooled, and concentrated before flash-freezing for storage at -80° C. The protocol for full-length MadS was similar, except that the final protein was stored in 50 mM Tris pH 8.0, 100 mM sodium chloride, 10% glycerol, 0.05% DDM (w/v). LiaR was purified as previously described [19]. MadR was purified similarly, except that the final protein was stored in 50 mM Tris pH 8.0, 100 mM sodium chloride, and 10% glycerol.

### Microscale thermophoresis (MST)

LiaS and MadS were labeled with the Red 2^nd^-generation NHS dye (Nanotemper, MO-L011) according to manufacturer instructions, aliquoted and stored at -80° C as needed. Before MST experiments were conducted, all proteins were spun at 20,000x g for 10 minutes at 4° C to pellet aggregates. Concentration and degree-of-labeling were determined by measuring absorbances at 280 nm (baseline 340 nm correction) and 650 nm, 20 nM of labeled LiaS or MadS were mixed 1:1 with serial dilutions of either MadR (180 µM to 5.49 nM) or LiaR (248 µM to 7.57 nM), with the exception that LiaR was used in a serial dilution from 67.2 µM to 2.05 nM for labeled MadS only. The dilutions were incubated for 100 minutes at room temperature in the dark. Microscale thermophoresis was performed using premium capillaries (Nanotemper, MO-K025) with medium IR power at an automatically determined excitation power (100% for LiaS*-LiaR, 80% for LiaS*-MadR, 40% for MadS*-MadR, 60% for MadS*-LiaR) on a Monolith X instrument (Nanotemper). Data was analyzed using the MO.Control software (Nanotemper).

## ACKNOWLEDGEMENTS

The authors would like to thank Rachael Whitehead for assistance with graphical design. WRM is supported by NIH/NIAID R21 AI175821 and R21 AI190338. CAA is supported by an NIH/National Institute of Allergy and Infectious Diseases (NIAID) grant number K24 AI121296, R01 AI148342, R01 AI134637, and P01 AI152999. KSH and CAA are supported by the Department of Defense PRMRP Award HT9425-25-1-005.

